# The caninized anti-canine PD-1 monoclonal antibody in canine oral malignant melanoma: Efficacy and exploratory biomarker analysis

**DOI:** 10.1101/2025.08.26.671889

**Authors:** Masaya Igase, Kenji Hagimori, Sakuya Inanaga, Hiroki Mizoguchi, Kazuhito Itamoto, Masashi Sakurai, Tomoki Motegi, Hiroka Yamamoto, Masahiro Kato, Toshinori Shiga, Toshihiro Tsukui, Tetsuya Kobayashi, Takuya Mizuno

**Affiliations:** Laboratory of Molecular Diagnostics and Therapeutics, Joint Faculty of Veterinary Medicine, Yamaguchi University, Yamaguchi, Japan; Research Institute for Cell Design Medical Science, Yamaguchi University, Yamaguchi, Japan; Japan Small Animal Cancer Center, Affiliated with the Japan Small Animal Medical Center Foundation, Tokorozawa, Saitama, Japan; Kyoto Animal Medical Center, Kyoto, Japan; Kamogawa Animal Medical Center, Kyoto, Japan; Laboratory of Veterinary Small Animal Clinical Science, Joint Faculty of Veterinary Medicine, Yamaguchi University, Yamaguchi, Japan; Laboratory of Veterinary Pathology, Joint Faculty of Veterinary Medicine, Yamaguchi University, Yamaguchi, Japan; Veterinary Clinical Genetics and Information, Cooperative Department of Veterinary Medicine, Tokyo University of Agriculture and Technology, Tokyo, Japan; Nippon Zenyaku Kogyo Co., Ltd., Fukushima, Japan

**Keywords:** biomarker, canine, melanoma, microsatellite instability, PD-1 blockade

## Abstract

**Background:** Canine oral malignant melanoma (OMM) is a highly aggressive tumor with few treatment options. Because of its biological similarity to human mucosal melanoma, canine OMM represents a valuable spontaneous model for translational immunotherapy studies. Anti-PD-1 antibody therapy has shown promise in canine OMM; however, predictive biomarkers for treatment response and survival have not been identified.

**Methods:** We conducted a multicenter, prospective, investigator-initiated clinical trial to evaluate the safety and efficacy of a caninized anti-canine PD-1 monoclonal antibody (ca-4F12-E6) in 150 dogs with advanced OMM. The dogs were administered 3 mg/kg ca-4F12-E6 intravenously every two weeks. Treatment response was assessed using cRECIST v1.0. Biomarker analyses included peripheral blood parameters, cytokine/chemokines, peripheral lymphocyte subsets, microsatellite instability (MSI), immunohistochemistry for immune cell and mismatch repair protein markers, and RNA sequencing of the tumor tissue. Associations with clinical outcomes were determined by logistic regression and Cox proportional hazard models.

**Results:** The best overall response rate was 16.7%. Treatment-related adverse events occurred in 40.0% of the dogs, which were primarily grade 1–3. Increased baseline white blood cell, neutrophil count, and C-reactive protein (CRP) levels were significantly associated with poor response, shorter progression-free survival (PFS), and reduced overall survival (OS). MSI-high tumors were associated with significantly prolonged OS compared with MSI-low/microsatellite stable tumors. Transcriptome analysis revealed differentially expressed genes and enriched immune-related pathways in responders, though limited by sample size.

**Conclusion:** ca-4F12-E6 exhibited durable antitumor activity and an acceptable safety profile in dogs with OMM. Baseline systemic inflammatory markers and MSI status may serve as predictive biomarkers for clinical outcomes. The results support the use of canine OMM as a comparative model for human immuno-oncology and biomarker discovery.

## Background

Malignant melanoma in dogs frequently originates from the oral mucosa and is recognized as a relevant spontaneous model for human mucosal melanoma [1]. Oral malignant melanoma (OMM) is the most common malignant oral tumor in dogs and is characterized by aggressive local invasion and a high rate of metastasis [1]. Although surgical excision and radiation therapy are the standard treatments used to achieve local control, the prognosis for advanced OMM remains poor, particularly because of the frequent recurrence and distant metastasis [2]. Conventional chemotherapy does not provide a meaningful improvement in survival [3]. These challenges highlight the urgent need for more effective systemic therapies to improve the clinical management of advanced OMM [4].

In humans, immune checkpoint inhibitors (ICIs) that target programmed cell death protein 1 (PD-1) or programmed cell death ligand 1 (PD-L1) have revolutionized the treatment of various malignancies, including melanoma [5,6]. By restoring antitumor immune responses, PD-1 blockade results in durable responses for patients with advanced melanoma [7]. Because of the biological and clinical similarities between canine OMM and human mucosal melanoma [1], the development and clinical evaluation of anti-PD-1 immunotherapy in dogs represent a promising treatment approach.

In contrast to human studies, the development of ICIs in veterinary fields is at an early stage. Since the initial report in 2017 by Maekawa et al. describing a canine chimeric anti-PD-L1 antibody (c4G12) [8], several anti-PD-1/PD-L1 antibodies have been developed. Of these, only the canine chimeric anti-PD-L1 antibody c4G12 and the anti-canine PD-1 antibodies [9] developed by our group, which includes a canine chimeric antibody (ch-4F12-E6) and a caninized antibody (ca-4F12-E6), have progressed to veterinary investigator-initiated clinical trials. An anti-canine PD-1 antibody (MP001) developed by Biocytogen Pharmaceuticals (Beijing, China) was described in a single case of oral adenocarcinoma [10]. In addition, a caninized anti-PD-1 antibody (gilvetmab) developed by Merck has received conditional approval from the USDA for canine tumors and has become commercially available, although detailed clinical data have not been published. Both c4G12 and ca-/ch-4F12-E6 prolong the survival time of dogs with stage 4 OMM compared with historical controls, with overall response rates (ORR) of approximately 15% [8,9]. Although the number of enrolled cases in these studies was relatively small, the limited number of responders highlights the necessity of identifying predictive biomarkers to optimize treatment selection.

Prognostic biomarker studies for PD-1/PD-L1 blockade have significantly advanced in humans. Predictive factors, such as tumor PD-L1 expression [11], tumor mutational burden (TMB) [12], mismatch repair deficiency (dMMR) [13], high microsatellite instability (MSI-high) [14], tumor-infiltrating lymphocyte density [15,16], interferon-gamma (IFN-γ)-related gene signatures [17], and peripheral blood markers, including neutrophil and lymphocyte counts and lactate dehydrogenase (LDH) levels, have been proposed [18]. However, because of the variability in cutoff values and predictive accuracy across tumor types and assay methods, combining multiple biomarkers is recommended for clinical decision-making. In canine patients, Maekawa et al. (2022) found that lower baseline serum levels of prostaglandin E2 (PGE2), monocyte chemoattractant protein-1 (MCP-1), and vascular endothelial growth factor-A (VEGF-A), as well as higher levels of interleukin (IL)-2, IL-12, and stem cell factor, were associated with prolonged OS in dogs administered c4G12 for OMM [19]; however, no biomarkers that predict response to caninized anti-PD-1 antibody therapy have been identified.

In this study, we conducted a multicenter, prospective, investigator-initiated clinical trial to evaluate the efficacy of the caninized anti-canine PD-1 antibody (ca-4F12-E6) in dogs with OMM. We also evaluated predictive biomarkers by performing genomic analysis and immunohistochemistry on tumor tissues, along with a comprehensive analysis of blood-based biomarkers.

## Materials and methods

### Ethics statement

The design of this clinical study involving dogs was approved by the Ethics Review Board of the Joint Faculty of Veterinary Medicine at Yamaguchi University (Approval No. 17) and the Ethics Review Board of the Kyoto Animal Medical Center (Approval No.2). The owners all agreed and provided written informed consent.

### Study design and treatments

This study was an open-label, nonrandomized, double-center clinical trial conducted in dogs diagnosed with oral malignant melanoma to evaluate the efficacy of the anti-PD-1 antibody and identify putative prognostic biomarkers. The study was performed at the Yamaguchi University Animal Medical Center (Yamaguchi, Japan) and the Kyoto Animal Medical Center (Kyoto, Japan) from July 20, 2020, to May 5, 2024. Inclusion criteria were as follows: (1) dogs diagnosed with OMM by cytology or histopathology, (2) tumors that were measurable by either caliper, X-ray, or computed tomography (CT) scan, (3) a life expectancy of at least one month, and (4) prior treatment with surgery, radiation, or chemotherapy were acceptable if the animal had progressive disease at the beginning of the trial. Cases were excluded if they were concurrently treated with systemic treatment, such as cytotoxic agents, at the beginning of the trial or if pet owners did not provide informed consent. All cases were restaged by X-ray or CT scan before initiation of the study.

The production of caninized anti-canine PD-1 antibody ca-4F12-E6 has been described [9]. Based on our previous studies [9,20], eligible cases received intravenous infusions of 3 mg/kg of ca-4F12-E6 every 2 weeks for each 10-week treatment cycle. Regardless of treatment response, continuous administration of ca-4F12-E6 was continued beyond the initial assessment period, only if acceptable adverse events (AEs) were observed, the dogs were assessed by the investigators to be in better condition compared with preenrollment, and the owner opted to continue the treatment.

### Safety and efficacy assessment

The assessment of AEs was graded according to the Veterinary Cooperative Oncology Group-Common Terminology Criteria for Adverse Events v2 (VCOG-CTCAE v2) scale [21]. Assessment was from the initial treatment to trial termination. Physical examinations and blood tests (complete blood count and blood chemistry) were conducted every two weeks to monitor AEs. Treatment-related AEs were defined by the VCOG-CTCAE v2, based on the investigator’s clinical judgment regarding the potential association with the treatment.

Tumor response was assessed according to the response evaluation criteria for solid tumors in dogs cRECIST v1.0 [22]. It was achieved using caliper, X-ray, or CT scans by comparing tumor diameters between the baseline and the end of each treatment cycle. A target lesion was defined as a tumor with a diameter greater than 10 mm, whereas a nontarget lesion included tumors with a diameter <10 mm. For cases in which the primary oral tumor could not be identified as a target lesion because of prior treatment, and lung metastases were present, pulmonary lesions measuring 5 mm or more on CT scans were considered target lesions. Tumor response was evaluated for both target and nontarget lesions based on cRESIST v1.0 criteria every 10 weeks, which corresponded to two weeks after the completion of one treatment cycle consisting of five doses, except for cases in which disease progression (PD) was observed during the treatment cycle, resulting from clinical deterioration. The occurrence of a new metastatic lesion, progression of nontarget lesions, or clinical deterioration was defined as PD, even if the target lesion showed regression. Cases in which the tumor diameter exhibited a change of less than a 20% increase or a 30% decrease following the completion of each treatment cycle (every 10 weeks) were categorized as stable disease (SD). A complete response (CR) was defined as the disappearance of all measurable tumor lesions, whereas a partial response (PR) was defined as a reduction of at least 30% in the sum of tumor lesion diameters. Cases in which no evaluation was available because of loss to follow-up were classified as not evaluable (NE). The ORR was calculated as the proportion of dogs achieving either a CR or PR at the end of the first treatment cycle. The clinical benefit rate (CBR) was defined as the proportion of dogs achieving an objective response (CR, PR, or SD). Overall survival (OS) was defined as the duration from the initiation of treatment to death from any cause. Progression-free survival (PFS) was defined as the time from the initiation of treatment to either PD or death at the end of the study.

### Exploratory biomarker analysis

To identify predictive biomarkers for treatment response and prognosis, we selected candidate markers based on previously reported studies involving humans [23,24]. Baseline whole blood, serum, and tumor tissues were collected from a subset of enrolled dogs with owner consent. Whole blood samples were subjected to flow cytometry analysis of CD4^+^ and CD8^+^ T cells, and Foxp3^+^ Tregs, and DNA was extracted for microsatellite (MS) genomic analysis. Serum samples were analyzed using a multiplex cytokine/chemokine array. DNA and RNA samples from fresh tumor tissues were used for MS genomic analysis and comprehensive gene expression analysis, respectively, whereas the remaining tissues were collected as a formalin-fixed, paraffin-embedded (FFPE) tissue sample and used for diagnosis and immunohistochemistry for immune markers.

### PCR for microsatellite sequences

Genomic DNA was extracted from paired samples of whole blood and tumor tissue using a QIAamp DNA Blood Mini Kit (QIAGEN Tokyo, Tokyo, Japan) or a QIAamp DNA Mini Kit (QIAGEN Tokyo), respectively. As described previously [25], 12 MS sequences were amplified from the extracted DNA. The PCR amplicons were analyzed by capillary electrophoresis (3500xL Genetic Analyzer, Thermo Fisher Scientific K.K., Tokyo, Japan) by the DNA Core Facility of the Center for Gene Research, Yamaguchi University, and using a software application (Peak Scanner, Thermo Fisher Scientific K.K.). For each MS sequence, electropherograms of tumor and normal DNA were compared and defined as MSI-positive if a difference in product size was detected. Based on our previous report [25], a tumor sample was defined as MSI-low if <2 sequences were mutated or MSI-high if ≥2 sequences were mutated. Tumor samples with no detectable MS alterations were considered microsatellite stable (MSS). The electropherograms for each sample were evaluated independently by three observers.

### Flow cytometry

Peripheral blood mononuclear cells (PBMCs) were isolated from whole blood by density gradient centrifugation using Lymphoprep (Axis-Shield, Oslo, Norway) as described previously [9]. The PBMCs were stained with anti-canine CD3-FITC/CD4-RPE/CD8-Alexa Fluor647 cocktail antibody (Bio-Rad Laboratories, Hercules, CA) or multiple IgG1/IgG2a/IgG1 negative control antibodies (Bio-Rad Laboratories) to determine the proportion of T-cell subsets. To assess the Treg proportion, PBMCs were fixed and permeabilized using the Foxp3/Transcription Factor Staining Buffer Set (Invitrogen, Waltham, MA), and stained with anti-canine CD4-FITC antibody (YKIX302.9, eBioscience, San Diego, CA) followed by anti-human Foxp3-PE antibody (FJK016s, eBioscience) as described previously [26]. The cells were subject to flow cytometry (CytoFLEX, Beckman Coulter, Brea, CA) and the data were analyzed using FlowJo software (Treestar, San Carlos, CA).

### Multiplex cytokine/chemokine array

Serum samples were evaluated for cytokine expression, including granulocyte-macrophage colony-stimulating factor (GM-CSF), IFN-γ, IL-2, IL-6, IL-7, IL-8, IL-15, IP-10, keratinocyte-derived chemokine (KC), IL-10, IL-18, monocyte chemotactic protein-1 (MCP-1), and tumor necrosis factor (TNF)-alpha using the MILLIPLEX Canine Cytokine/Chemokine Magnetic Bead Panel (Merck Millipore, Burlington, MA, USA).

### Comprehensive RNA sequencing analysis

Fresh tumor tissues obtained by excision biopsy were preserved in a sample solution for DNA/RNA (Takara-Bio, Shiga, Japan) to prevent RNA degradation. Briefly, total RNA was purified from tumor tissue using the RNeasy Plus Mini Kit (QIAGEN, Hilden, Germany) based on the manufacturer’s protocol. Whole transcriptome analysis using the Illumina NovaSeq 6,000 sequencing system (Illumina, San Diego, CA) was outsourced to Novogene (Beijing, China). Paired-end 150 bp reads were collected, quality checked, and the adapter sequences were trimmed using Trim Galore ver. 0.6.3. Canine genome assembly (CanFam3.1) was achieved using STAR v2.7.3a, and transcript abundance was quantified using RSEM v1.3.3(21816040) with Ensembl gene transfer files (CanFam3.1.98, https://may2021.archive.ensembl.org/Canis_lupus_familiaris/). The raw expression count data were normalized using the multi-step trimmed mean of the M values method, and differentially expressed genes (DEGs) between CR+PR and SD+PD cases were identified using the estimateDE function in EdgeR (TCC v1.32.0) [27] in R (v4.2.1) and extracted based on a false discovery rate threshold of <0.01. Because the number of DEGs between the groups was limited, we concluded that conventional GSEA was insufficient to detect pathway-level changes. Therefore, we applied single-sample GSEA (ssGSEA) to evaluate enrichment scores for each sample, which enables pathway activity estimation, even in the absence of a clear set of DEGs [28]. After filtering out low-expressed genes, Ensembl gene IDs were converted to HGNC symbols to match the gene sets. ssGSEA was performed with the GSVA R package (v1.52.3), using immunologic signature gene sets from the Molecular Signatures Database (MSigDB v7.5.1), to quantify signature change scores for each sample. Differences in signature scores between groups were assessed using the Wilcoxon test.

### Immunohistochemistry

Infiltration of immune cells into tumors, including CD3^+^ T cells, CD8^+^ T cells, CD20^+^ B cells, Foxp3^+^ Treg cells, and CD204^+^ macrophages, was assessed by IHC, as previously described [20,29]. To examine DNA mismatch repair deficiency (dMMR) in the canine tumor samples, immunolabeling of MMR proteins, such as MSH2, MSH6, and MLH1, was performed as described previously [30]. Briefly, tumor samples were incubated with 10% H_2_O_2_ in phosphate-buffered saline for 10 min at 65°C for melanin bleaching. The sections were stained with rat anti-human CD3 monoclonal antibody (clone CD3-12, Abcam, Tokyo, Japan); rat anti-canine CD8 monoclonal antibody (clone F3-B2, own production [31]); rabbit anti-human CD20 polyclonal antibody (Thermo Fisher Scientific, Waltham, MA); rat anti-mouse FOXP3 monoclonal antibody (clone FJK-16s, Invitrogen, Carlsbad, CA); mouse anti-human MSR-A/CD204 monoclonal antibody (clone SRA-E5, Cosmo Bio Co., Ltd., Tokyo, Japan); mouse anti-human MLH1 monoclonal antibody (clone G168-15, GeneTex, Irvine, CA); rabbit anti-human MSH2 monoclonal antibody (clone SP46, Abcam); and mouse anti-human PMS2 monoclonal antibody (clone 163C1251, Novus Biologicals) as the primary antibodies. Histofine Simple Stain MAX PO Kit (Nichirei Corporation, Tokyo, Japan) was used as the secondary antibody. Histofine Simple Stain AEC (Nichirei Corporation, Tokyo, Japan) was used for visualization, and the slides were counterstained with Mayer’s hematoxylin (Wako, Osaka, Japan). For the immune cell markers, positive cells were manually counted in 10 randomly selected hotspots under high-power magnification (×400 HPF) for each sample, as described in a previous report [32]. Immunoreactivity was considered positive if the cell nuclei within tumor cells were immunolabeled with each anti-MMR antibody compared with isotype controls [30]. A veterinary pathologist reviewed and examined all samples to confirm and validate staining quality with all of the antibodies.

### Endpoints

The primary endpoint was ORR, and the secondary endpoints were safety, CBR, PFS, and OS. The exploratory objective was to identify genetic, pathological, immunological, or clinical predictive biomarkers associated with response and prognosis.

### Statistical analysis

Safety and efficacy were evaluated in all cases that received at least one treatment dose. Treatment-related AEs was recorded based on the physician’s clinical judgment, excluding neoplastic, infectious, metabolic, toxic, or other etiologies where possible. Univariate Cox proportional hazards models were used to evaluate the relationship between PFS or OS and baseline characteristics, as well as various biomarker parameters, including blood test results, multiplex cytokine/chemokine assays, PBMC subsets, MSI DNA testing, IHC for MMR markers, IHC for immune cell markers, and RNA sequencing data, as part of the exploratory objectives. In addition, logistic regression analysis was used to determine the relationship between the objective response and baseline characteristics, as well as biomarker parameters (as above). Kaplan–Meier analysis was used for a visual evaluation of PFS/OS stratified by MSI-high or MSI-low/MSS, followed by a Log-rank test. All analyses were performed using SAS (version 9.4 or later; SAS Institute Inc., Cary, NC, USA) or JMP Pro18.0.2 (JMP Statistical Discovery LLC, Cary, NC) software.

## Results

### Patients

We enrolled 150 dogs with OMM from two veterinary medical centers, which included 101 and 49 dogs from each institution, respectively (Table S1). The baseline characteristics are listed in Table 1. There were no significant differences in age, sex, breed, or tumor staging between the two cohorts. The median age of the dogs was 13 years. Among the breeds, Miniature dachshunds (28%) and Toy poodles (20%) were the most common. Based on the WHO staging system, 93 cases were classified as stage 4, 39 as stage 3, 15 as stage 2, and 3 as stage 1. Of the enrolled cases, 105 had a target lesion in the oral cavity as the primary site, whereas 45 had nonoral primary sites, such as lung metastasis and lymph node metastasis. Surgical resection was the most common prior treatment, which was performed in 46% of the cases. Figure 1 summarizes the treatment duration, treatment response, and the availability of various biological samples used for biomarker analysis in each case. As of the data cutoff date (May 5, 2024), the trial was ongoing in nine dogs (Figure 1A, B).

**Figure 1.**
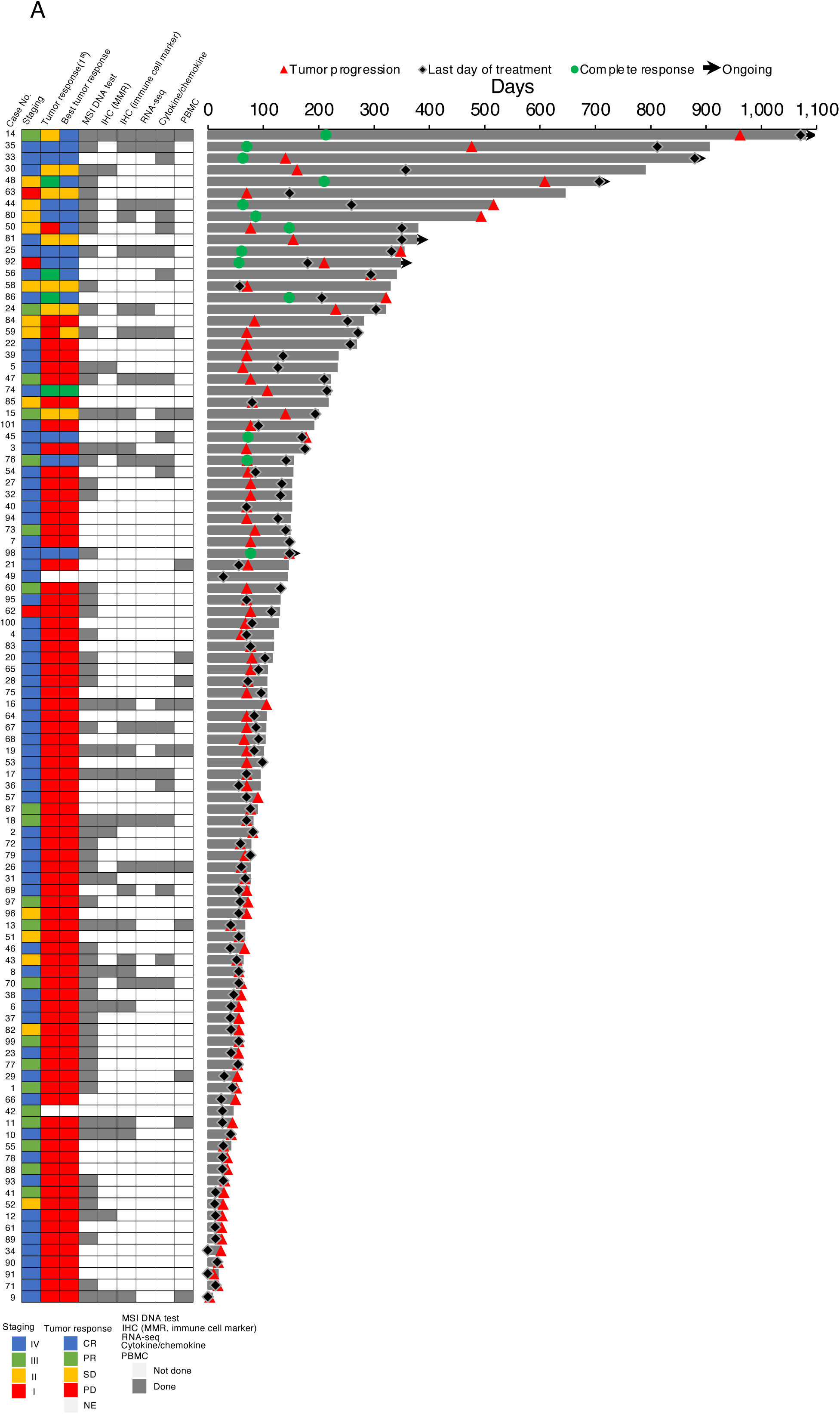
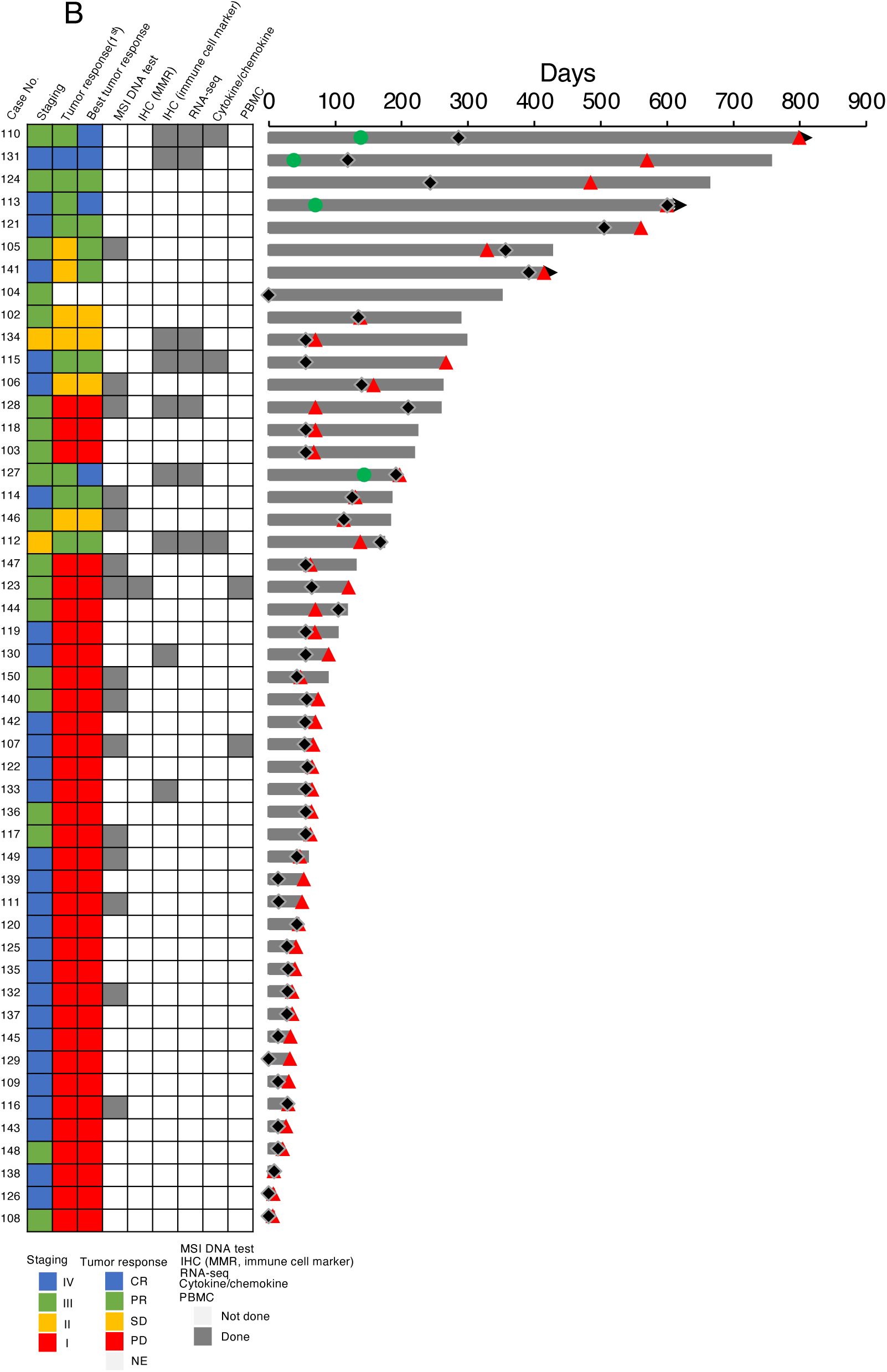
Overview of the clinical data and sample availability (A) Clinical timeline and sample collection summary for 101 dogs enrolled in the anti-PD-1 antibody clinical trial at Yamaguchi University. (B) Same overview as (A), shown for an additional 49 dogs at the Kyoto Animal Medical Center. The treatment duration, WHO staging, response evaluation, and availability of samples for various assays (MSI testing, IHC for MMR and immune cell markers, RNA-seq, cytokine/chemokine profiling, and PBMC isolation) are shown for each case. Treatment response categories include CR: complete response, PR: partial response, SD: stable disease, PD: progressive disease, and NE: not evaluable. The leftmost number beside each bar represents the individual case number and corresponds to the Case No. listed in Table S1.

**Table 1.**
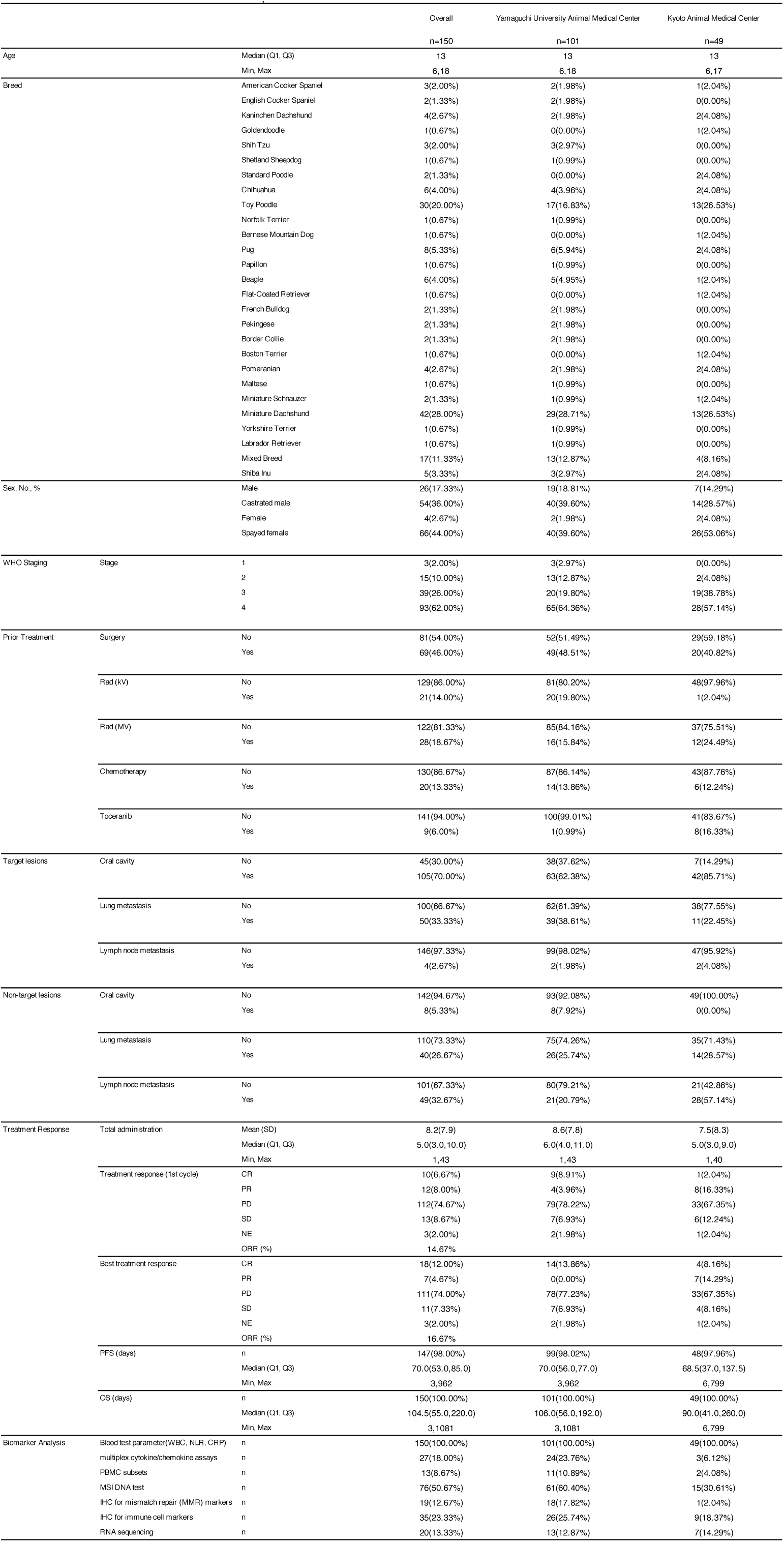
Baseline characteristsics and treatment response.

|  |  |  | Overall | Yamaguchi University Animal Medical Center | Kyoto Animal Medical Center |
| --- | --- | --- | --- | --- | --- |
|  |  |  | n=150 | n=101 | n=49 |
| Age | Median (Q1, Q3) |  | 13 | 13 | 13 |
|  | Min, Max |  | 6, 18 | 6, 18 | 6, 17 |
| Breed | American Cocker Spaniel |  | 3(2.00%) | 2(1.98%) | 1(2.04%) |
|  | English Cocker Spaniel |  | 2(1.33%) | 2(1.98%) | 0(0.00%) |
|  | Karinhchen Dackshund |  | 4(2.67%) | 2(1.98%) | 2(4.08%) |
|  | Goldendoodle |  | 1(0.67%) | 0(0.00%) | 1(2.04%) |
|  | Shih Tzu |  | 3(2.00%) | 3(2.97%) | 0(0.00%) |
|  | Shetland Sheepdog |  | 1(0.67%) | 1(0.99%) | 0(0.00%) |
|  | Standard Poodle |  | 2(1.33%) | 0(0.00%) | 2(4.08%) |
|  | Chihuahua |  | 6(4.00%) | 4(3.96%) | 2(4.08%) |
|  | Toy Poodle |  | 30(20.00%) | 17(16.83%) | 13(26.53%) |
|  | Norfolk Terrier |  | 1(0.67%) | 1(0.99%) | 0(0.00%) |
|  | Bernese Mountain Dog |  | 1(0.67%) | 0(0.00%) | 1(2.04%) |
|  | Pug |  | 8(5.33%) | 6(5.94%) | 2(4.08%) |
|  | Papillon |  | 1(0.67%) | 1(0.99%) | 0(0.00%) |
|  | Beagle |  | 6(4.00%) | 5(4.95%) | 1(2.04%) |
|  | Flat-Coated Retriever |  | 1(0.67%) | 0(0.00%) | 1(2.04%) |
|  | French Bulldog |  | 2(1.33%) | 2(1.98%) | 0(0.00%) |
|  | Pekingese |  | 2(1.33%) | 2(1.98%) | 0(0.00%) |
|  | Border Collie |  | 2(1.33%) | 2(1.98%) | 0(0.00%) |
|  | Boston Terrier |  | 1(0.67%) | 0(0.00%) | 1(2.04%) |
|  | Pomeranian |  | 4(2.67%) | 2(1.98%) | 2(4.08%) |
|  | Maltese |  | 1(0.67%) | 1(0.99%) | 0(0.00%) |
|  | Miniature Schnauzer |  | 2(1.33%) | 1(0.99%) | 1(2.04%) |
|  | Miniature Dachshund |  | 42(28.00%) | 29(28.71%) | 13(26.53%) |
|  | Yorkshire Terrier |  | 1(0.67%) | 1(0.99%) | 0(0.00%) |
|  | Labrador Retriever |  | 1(0.67%) | 1(0.99%) | 0(0.00%) |
|  | Mixed Breed |  | 17(11.33%) | 13(12.87%) | 4(8.16%) |
|  | Shiba Inu |  | 5(3.33%) | 3(2.97%) | 2(4.08%) |
| Sex, No., % | Male |  | 26(17.33%) | 19(18.81%) | 7(14.29%) |
|  | Castrated male |  | 54(36.00%) | 40(39.60%) | 14(28.57%) |
|  | Female |  | 4(2.67%) | 2(1.98%) | 2(4.08%) |
|  | Spayed female |  | 66(44.00%) | 40(39.60%) | 26(53.06%) |
| WHO Staging | Stage | 1 | 3(2.00%) | 3(2.97%) | 0(0.00%) |
|  |  | 2 | 15(10.00%) | 13(12.87%) | 2(4.08%) |
|  |  | 3 | 39(26.00%) | 20(19.80%) | 19(38.78%) |
|  |  | 4 | 93(62.00%) | 65(64.36%) | 28(57.14%) |
| Prior Treatment | Surgery | No | 81(54.00%) | 52(51.49%) | 29(59.18%) |
|  |  | Yes | 69(46.00%) | 49(48.51%) | 20(40.82%) |
|  | Rad (kV) | No | 129(86.00%) | 81(80.20%) | 48(97.96%) |
|  |  | Yes | 21(14.00%) | 20(19.80%) | 1(2.04%) |
|  | Rad (MV) | No | 122(81.33%) | 85(84.16%) | 37(75.51%) |
|  |  | Yes | 28(18.67%) | 16(15.84%) | 12(24.49%) |
|  | Chemotherapy | No | 130(86.67%) | 87(86.14%) | 43(87.76%) |
|  |  | Yes | 20(13.33%) | 14(13.86%) | 6(12.24%) |
|  | Toceranib | No | 141(94.00%) | 100(99.01%) | 41(83.67%) |
|  |  | Yes | 9(6.00%) | 1(0.99%) | 8(16.33%) |
| Target lesions | Oral cavity | No | 45(30.00%) | 38(37.62%) | 7(14.29%) |
|  |  | Yes | 105(70.00%) | 63(62.38%) | 42(85.71%) |
|  | Lung metastasis | No | 100(66.67%) | 62(61.39%) | 38(77.55%) |
|  |  | Yes | 50(33.33%) | 39(38.61%) | 11(22.45%) |
|  | Lymph node metastasis | No | 146(97.33%) | 99(98.02%) | 47(95.92%) |
|  |  | Yes | 4(2.67%) | 2(1.98%) | 2(4.08%) |
| Non-target lesions | Oral cavity | No | 142(94.67%) | 93(92.08%) | 49(100.00%) |
|  |  | Yes | 8(5.33%) | 8(7.92%) | 0(0.00%) |
|  | Lung metastasis | No | 110(73.33%) | 75(74.26%) | 35(71.43%) |
|  |  | Yes | 40(26.67%) | 26(25.74%) | 14(28.57%) |
|  | Lymph node metastasis | No | 101(67.33%) | 80(79.21%) | 21(42.86%) |
|  |  | Yes | 49(32.67%) | 21(20.79%) | 28(57.14%) |
| Treatment Response | Total administration | Mean (SD) | 8.2(7.9) | 8.6(7.8) | 7.5(8.3) |
|  |  | Median (Q1, Q3) | 5.0(3.0, 10.0) | 6.0(4.0, 11.0) | 5.0(3.0, 9.0) |
|  |  | Min, Max | 1, 43 | 1, 43 | 1, 40 |
|  | Treatment response (1st cycle) | CR | 10(6.67%) | 9(8.91%) | 1(2.04%) |
|  |  | PR | 12(8.00%) | 4(3.96%) | 8(16.33%) |
|  |  | PD | 112(74.67%) | 79(78.22%) | 33(67.35%) |
|  |  | SD | 13(8.67%) | 7(6.93%) | 6(12.24%) |
|  |  | NE | 3(2.00%) | 2(1.98%) | 1(2.04%) |
|  |  | ORR (%) | 14.67% |  |  |
|  | Best treatment response | CR | 18(12.00%) | 14(13.86%) | 4(8.16%) |
|  |  | PR | 7(4.67%) | 0(0.00%) | 7(14.29%) |
|  |  | PD | 111(74.00%) | 78(77.23%) | 33(67.35%) |
|  |  | SD | 11(7.33%) | 7(6.93%) | 4(8.16%) |
|  |  | NE | 3(2.00%) | 2(1.98%) | 1(2.04%) |
|  |  | ORR (%) | 16.67% |  |  |
|  | PFS (days) | n | 147(98.00%) | 99(98.02%) | 48(97.96%) |
|  |  | Median (Q1, Q3) | 70.0(53.0, 85.0) | 70.0(56.0, 77.0) | 68.5(37.0, 137.5) |
|  |  | Min, Max | 3, 962 | 3, 962 | 6, 799 |
|  | OS (days) | n | 150(100.00%) | 101(100.00%) | 49(100.00%) |
|  |  | Median (Q1, Q3) | 104.5(55.0, 220.0) | 106.0(56.0, 192.0) | 90.0(41.0, 260.0) |
|  |  | Min, Max | 3, 1081 | 3, 1081 | 6, 799 |
| Biomarker Analysis | Blood test parameter(WBC, NLR, CRP) | n | 150(100.00%) | 101(100.00%) | 49(100.00%) |
|  | multiplex cytokine/chemokine assays | n | 27(18.00%) | 24(23.76%) | 3(6.12%) |
|  | PBMC subsets | n | 13(8.67%) | 11(10.89%) | 2(4.08%) |
|  | MSI DNA test | n | 76(50.67%) | 61(60.40%) | 15(30.61%) |
|  | IHC for mismatch repair (MMR) markers | n | 19(12.67%) | 18(17.82%) | 1(2.04%) |
|  | IHC for immune cell markers | n | 35(23.33%) | 26(25.74%) | 9(18.37%) |
|  | RNA sequencing | n | 20(13.33%) | 13(12.87%) | 7(14.29%) |

### Treatment-related adverse events

All 150 enrolled dogs were evaluated for safety, and they received a median of 5 doses (range, 1–43) of anti-PD-1 antibody during the study period. The treatment-related AEs are listed in Table 2. Treatment-related AEs occurred in 60 dogs (40.00%). Most AEs were mild to moderate in severity (Grade ≤3). The most frequently observed AE was diarrhea, which occurred in 14.00% of the cases. Only one dog experienced a Grade 4 elevation in bilirubin levels associated with Grade 3 pancreatitis, which resolved following the cessation of antibody administration. Immune-related AEs included pruritus (n = 3, 2.00%), hypopigmentation (n = 1, 0.67%), neutropenia (n = 1, 0.67%), and immune-mediated hemolytic anemia (IMHA, n = 1, 0.67%). No treatment-related Grade 5 events were observed.

**Table 2.** Treatment-related adverse events.

|  | Adverse events, all grades, n |  |  |  |  |
| --- | --- | --- | --- | --- | --- |
|  | Grade 1 | Grade 2 | Grade 3 | Grade 4 | Total (%) |
| Appetite, altered | 11 | 0 | 1 | 0 | 12 (8.0) |
| Anorexia | 4 | 0 | 0 | 0 | 4 (2.7) |
| Lethargy/Fatigue | 10 | 0 | 0 | 0 | 10 (6.7) |
| Diarrhoea | 14 | 7 | 0 | 0 | 21 (14) |
| Vomiting | 8 | 1 | 0 | 0 | 9 (6.9) |
| Colitis | 0 | 2 | 0 | 0 | 2 (1.3) |
| ALT | 3 | 5 | 7 | 0 | 15 (10) |
| AST | 5 | 1 | 2 | 0 | 8 (5.3) |
| ALP | 6 | 1 | 2 | 0 | 9 (6.0) |
| T.Bil | 0 | 0 | 0 | 1 | 1(0.7) |
| BUN | 1 | 6 | 0 | 0 | 7 (4.7) |
| Cre | 2 | 0 | 1 | 0 | 3 (2.0) |
| Pruritus | 3 | 0 | 0 | 0 | 3 (2.0) |
| Hypopigmentation<br>/vitiligo | 0 | 1 | 0 | 0 | 1(0.7) |
| Neutropenia | 0 | 0 | 1 | 0 | 1(0.7) |
| IMHA, hemolysis | 0 | 0 | 1 | 0 | 1(0.7) |
| Pancreatis | 0 | 0 | 1 | 0 | 1(0.7) |

ALT, alanin transferase; AST, asparate aminotransferase; ALP, alkaline phosphatase;
T.Bil, total bilirubin; BUN, blood urea nitrogen; IMHA, immune-mediated hemolytic anemia

### Efficacy

Treatment response was assessed in all 150 enrolled dogs, including those with clinical PD, in which radiographic measurement of tumor size was not feasible, but disease progression was evident. After the first treatment cycle, 10 dogs (6.67%) achieved a CR, 12 (8.00%) achieved a PR, 11 (7.33%) had SD, 113 (75.33%) exhibited PD, and three (2.00%) were classified as NE (Table 1 and Figure 1). No significant differences in treatment response were observed between the two institutions. The ORR following the first treatment cycle was 14.67%. The CBR was 22.00%. Among the 95 dogs for whom tumor burden was evaluable after each treatment cycle, the maximum change from baseline in tumor burden is shown in Supplementary Figures 1a and 1b. Notably, 4 of these dogs (No. 5, 7, 68, and 69) showed a dissociated response, characterized by a reduction in the size of the target lesions, while showing progression in nontarget lesions, indicating heterogeneous responses across all tumor sites. Furthermore, four dogs (No. 14, 50, 105, and 141), who were initially classified as nonresponders, had delayed clinical responses with continued treatment (Supplementary Figure 1b). Therefore, considering the best responses observed during the entire study period, the ORR increased to 16.67% (Table 1).

The median PFS for 147 evaluable dogs was 70 days (range: 3–962), whereas the median OS for all 150 dogs was 105 days (range: 3–1,081). The median survival time for responders (CR + PR) and nonresponders (SD + PD) was 348 days and 68 days for PFS, and 516 days and 100 days for OS, respectively (Supplementary Figures 2a, b).

### Clinical response and biomarkers

Tumor staging and blood test parameters, including white blood cell (WBC) count, neutrophil count, neutrophil-to-lymphocyte ratio (NLR), and C-reactive protein (CRP), were determined for all cases. However, because of factors, such as tumor location, difficulty of performing anesthesia, and costs, procuring all types of biomarker samples from every case was not feasible. Other biomarker analyses could only be performed in a subset of cases (Figure 1A, B): microsatellite instability (MSI) testing (n = 76), IHC for MMR markers (n = 19), IHC for immune cell markers (n = 35), RNA sequencing (n = 20), multiplex cytokine/chemokine assays (n = 27), and PBMC subset analysis by flow cytometry (n = 13).

We performed a logistic regression analysis using the objective response (CR + PR) as the event, and the results are summarized in Supplementary Table 2. Although lymphocyte count and NLR were not significantly associated with treatment response, increased WBC count [odds ratio (OR): 0.856; 95% confidence interval (CI): 0.766– 0.957; p = 0.006], neutrophil count (OR: 0.829; 95% CI: 0.723–0.950; p = 0.007), and CRP level (OR: 0.716; 95% CI: 0.545–0.941; p = 0.016) were significantly associated with poor treatment response.

Other biomarker analyses did not have predictive utility. DEGs between responders (CR + PR) and nonresponders (SD + PD) were identified using a cutoff false discovery rate (FDR, q-value) <0.1 (Supplementary Table 3). Four genes showed significant changes in expression, including *VGLL2* (logFC = −8.702, q = 0.048), *TGM3* (logFC = 5.151, q = 0.048), *SLC24A5* (logFC = 3.991, q = 0.048), and *SLURP1* (logFC = 9.973, q = 0.057).

In addition, ssGSEA based on RNA-seq data (see in Supplementary Table 5) revealed considerable enrichment in genes associated with antitumor immunity, such as GSE36392_TYPE_2_MYELOID_VS_EOSINOPHIL_IL25_TREATED_LUNG_UP (OR:<0.001; 95% CI: <0.001–0.007; p = 0.041) and GSE3691_IFN_PRODUCING_KILLER_DC_VS_CONVENTIONAL_DC_SPLEEN_UP (OR:<0.001; 95% CI: <0.001–0.024; p = 0.040).

### Survival and biomarkers

Next, we conducted Cox proportional hazards model analyses using PFS and OS, and the results are summarized in Table 3 and Supplementary Table 4. Cases that achieved an objective response (CR + PR) exhibited significantly prolonged PFS (hazard ratio (HR): 0.183; 95% CI: 0.105–0.318; p < 0.001) and OS (HR: 0.190; 95% CI: 0.103–0.354; p < 0.001). Consistent with treatment response analysis, elevated baseline WBC counts, neutrophil counts, and CRP levels were significantly associated with shorter PFS and OS. Moreover, a higher peripheral lymphocyte count was significantly associated with a shorter PFS (HR: 1.165; 95% CI: 1.001–1.355; p = 0.048).

**Table 3.**
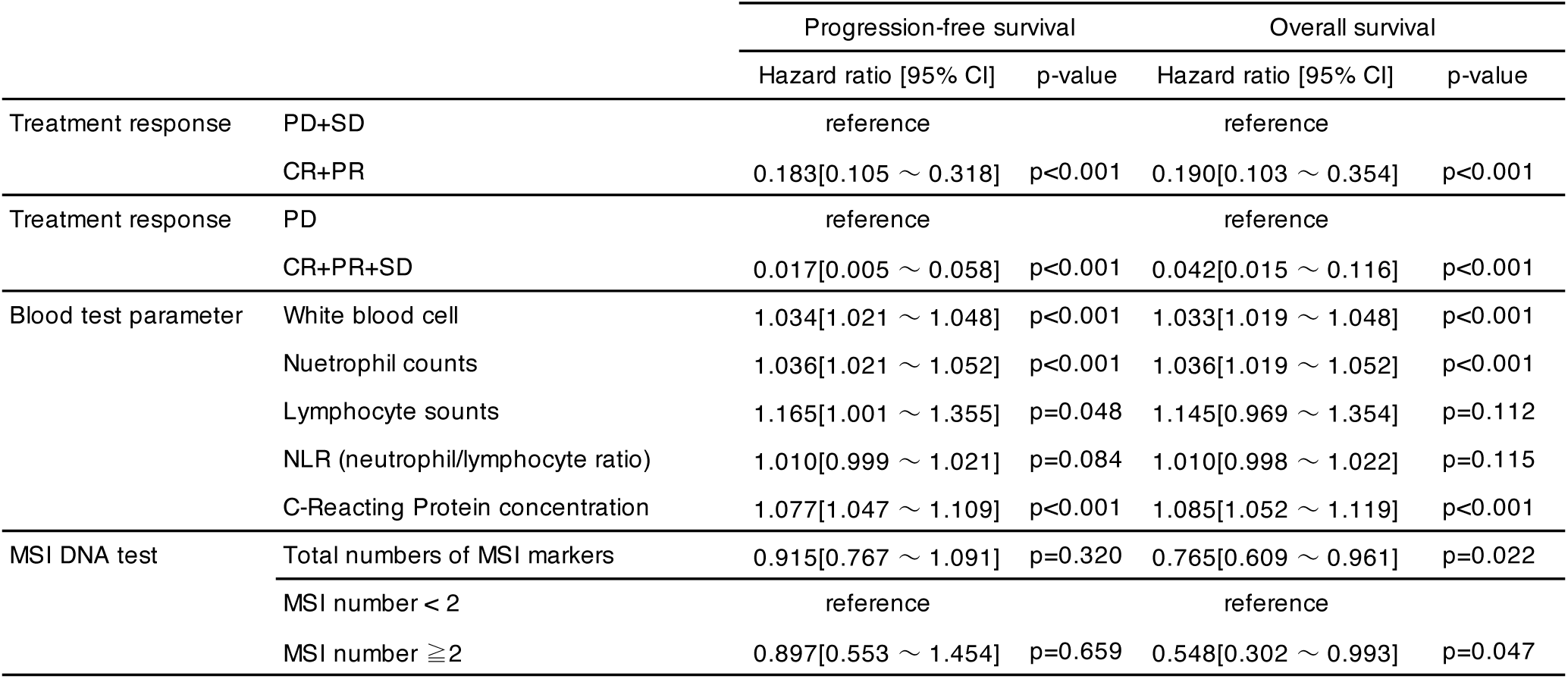
Cox proportional hazards model for prognosis.

With respect to the MSI DNA test, more MSI-positive loci were significantly associated with prolonged OS (HR: 0.765; 95% CI: 0.609–0.961; p = 0.022), but not with PFS (p = 0.320). Moreover, applying a cutoff of ≥2 MSI-positive loci to define MSI-high, which is indicative of dMMR [27], resulted in a significant association with prolonged overall survival (HR: 0.548; 95% CI: 0.302–0.993; p = 0.047). We classified cases with ≥2 MSI-positive loci as MSI-high and those with fewer loci as MSI-low/MSS. Using a Kaplan– Meier analysis for PFS and OS, the MSI-high cases exhibited a significantly longer OS compared with MSI-low/MSS cases (median OS: 200 days vs. 95 days), although no significant difference was observed in PFS (Figure 2).

**Figure 2.**
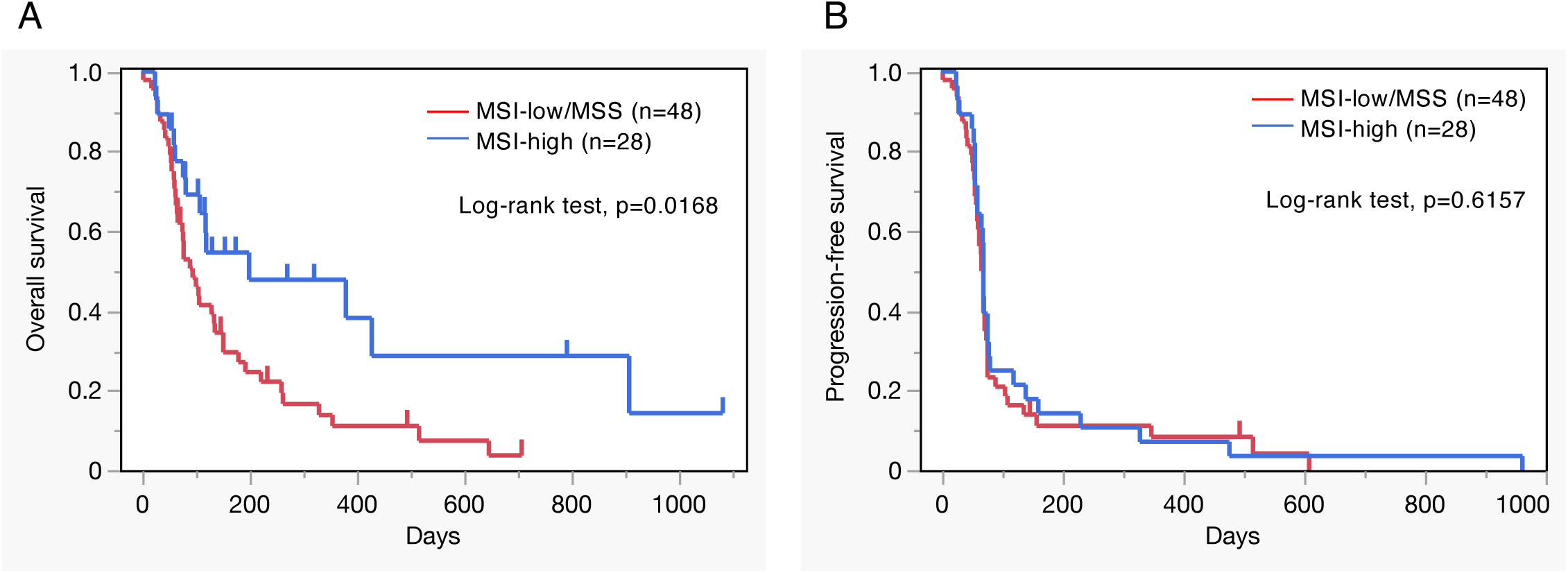
Survival analysis according to the microsatellite instability status (A) Kaplan–Meier curve for overall survival (OS) comparing microsatellite instability (MSI)-high (n = 28) and MSI-low/microsatellite stable (MSS) (n = 48) dogs. MSI-high status is associated with a significantly prolonged OS (log-rank test, p = 0.0168). (B) Kaplan–Meier curve for progression-free survival (PFS) in the same cohorts. No significant difference in PFS was observed (log-rank test, p = 0.6157).

For ssGSEA based on RNA-seq data (Supplementary Table 5), interpretation was limited because of the small number of available cases, which resulted in extensive confidence intervals and unstable hazard ratio estimates. However, the gene set GSE43955_1H_VS_42H_ACT_CD4_TCELL_WITH_TGFB_IL6_DN was significantly associated with prolonged PFS (HR <0.001; 95% CI: <0.001–0.052; p = 0.040). This suggests that downregulated genes associated with CD4+ T cell activation in the presence of TGF-β and IL-6 may portend a favorable prognosis.

## Discussion

This multicenter, prospective clinical trial is the largest to date, involving 150 enrolled dogs, to assess the efficacy and safety of a caninized anti-PD-1 monoclonal antibody (ca-4F12-E6) in dogs with OMM. We demonstrated a best ORR of 16.67%, which corroborates earlier single-institution studies using either c4G12 (anti-canine PD-L1 antibody, [8]) or the ca-4F12-E6 antibody [9], both of which achieved ORRs of ∼15% in far smaller cohorts. These results confirm that ICIs can achieve durable tumor control in a clinically meaningful subset of dogs with advanced OMM. In human mucosal melanoma, several clinical trials have reported an ORR of approximately 20% with PD-1 monotherapy, which is lower compared with the response rate of approximately 40% observed in cutaneous melanoma [33,34], which suggests a poorer response to ICIs. Therefore, novel therapeutic strategies, such as combination therapies, are highly anticipated in humans, which further enhances the value of the dog model as a translational platform for human mucosal melanoma.

Treatment-related AEs occurred in 40% of the dogs, with the majority being grade 1–3 gastrointestinal events. These results are comparable with those reported in previous clinical trials of ch-/ca-4F12-E6 and c4G12, which showed similar rates ranging from 35.1% to 63.3% [8,9,20,35,36]. Immune-related AEs, such as pruritus, hypopigmentation, neutropenia, and immune-mediated hemolysis, were rare (each observed in less than 3.0% of dogs), whereas no grade 5 AEs were reported. These results are consistent with the safety profiles reported for c4G12 [8,35,36], and support the safety of PD-1/PD-L1 blockade in veterinary medicine. Consistent with previous studies [9,20], the occurrence of a single grade 4 hepatobiliary AE serves as a reminder to clinicians that although severe toxicity is rare, close monitoring and prompt intervention are essential.

In four cases (No. 5, 7, 68, and 69), tumor shrinkage was observed in target lesions; however, the progression of nontarget lesions, including new lesions, was also evident. As a result, these cases were classified as PD under the current cRECIST criteria. In addition, four dogs (No. 14, 50, 105, and 141) who were initially categorized as nonresponders demonstrated clinical responses during later treatment cycles. These atypical response patterns resemble “dissociated response” or “late response/pseudoprogression,” which occur in a subset of patients with 7.5%–12.5% receiving ICIs in humans [37–39]. The results suggest that the antitumor effects of anti-PD-1 antibodies may take time to appear, and response rate can vary between different tumor sites. Unlike conventional chemotherapy, these results demonstrate that the timing of evaluation and the selection of target lesions can significantly impact the classification of treatment response. In humans, similar issues have resulted in the development of new response criteria, such as irRECIST [40] and iRECIST [41], to better reflect the unique patterns observed with ICIs. Under these criteria, disease progression identified at the first evaluation is considered unconfirmed PD, and treatment is continued until a second assessment confirms PD. Therefore, we suggest that updated response criteria are also needed in veterinary medicine to improve the current cRECIST criteria.

Systemic inflammation occurs in human cancer patients through the release of tumor-derived inflammatory cytokines, such as IL-6 and TNF-α, which suppress antitumor immunity and reduce the efficacy of PD-1 blockade [42,43]. Moreover, large-scale clinical datasets in humans have identified elevated absolute neutrophil counts and CRP levels as strong negative prognostic markers in patients administered ICIs [23,44]. In the present study, baseline leukocytosis, neutrophilia, and elevated CRP were associated with a poor response and shorter PFS and OS; however, based on the cytokine/chemokine multiplex assay, we did not observe significant differences in key cytokines, such as IL-6 or TNF-α, between responders and nonresponders. This may be the result of a limited sample size or the enrolled dogs displaying highly variable overall health conditions and stages; thus, further studies on a larger cohort are warranted. Nevertheless, complete blood counts and CRP are routinely measured in veterinary settings, making them practical and accessible biomarkers for predicting immunotherapy efficacy. These results suggest that systemic inflammatory markers can inform clinical decision-making in canine tumor cases.

Tumors with dMMR, which occur in <4% of human cancers, show high MSI because of a defect in one of the MMR proteins (e.g., MLH1, MSH2, MSH6, or PMS2) [45]. These tumors accumulate somatic mutations because of repair errors during DNA replication, resulting in a higher neoantigen load. Neoantigens are presented on tumor cells by MHC molecules and are recognized by CD8⁺ T cells, thereby enhancing the therapeutic efficacy of ICIs [46]. Of the 76 dogs evaluated for MSI, 28 (36.84%) were classified as MSI-high/dMMR based on the criteria established in our previous study [25]. MSI-H status occurs at a high frequency in human uterine corpus endometrial carcinoma (TCGA-UCEC; 31.37%), but at much lower rates in skin melanoma (TCGA-SKCM; 0.64%) and mucosal melanoma (Japanese cohort; 1.28%) [45,47]. These results suggest that canine OMM serves as a potential translational model for MSI-H tumors in humans. MSI-H tumors are frequently associated with high TMB, which enhances their responsiveness to ICIs [48]. Consistent with this finding, a large-scale genomic study of naturally occurring canine tumors found that OMM exhibits a relatively high TMB among various tumor types, which is comparable to that observed in osteosarcoma and hemangiosarcoma. It also revealed that the absolute TMB levels in dogs tend to be lower compared with those observed in human cancers [49]. Therefore, validating our MSI findings by calculating TMB through whole-exome sequencing in canine OMM will be an important step in future translational studies. Nonetheless, canine OMM with MSI-high exhibited significantly prolonged OS compared with MSI-low/MSS. This indicates that MSI status may serve as a predictive biomarker for the efficacy of anti-canine PD-1 antibody therapy.

In humans, comprehensive gene expression profiling using RNA-seq or single-cell RNA-seq, combined with IHC analysis of immune cell infiltration, has been used to identify biomarkers associated with antitumor immune responses [24]. Key features, such as an IFN-γ signature, expression of immune checkpoint molecules, and immune cell infiltration, have been linked to ICI-treatment outcomes. Notably, tumors with abundant CD8^+^T cell infiltration commonly referred to as “hot tumors,” tend to show better responses to ICIs. However, in canine OMM, no significant correlation was observed between T cell infiltration measured by IHC and treatment response. Our RNA-seq analysis identified four genes (VGLL2, TGM3, SLC24A5, and SLURP1) that were associated with a response to anti-canine PD-1 antibody. To date, these genes have not been associated with antitumor immunity, suggesting a novel finding specific to canine OMM. SLURP1 is known to suppress neutrophil adhesion to endothelial cells and migration into peripheral tissues [50]. Therefore, the observed association between SLURP1 upregulation and treatment response may be linked to the poorer outcomes observed in dogs with elevated absolute neutrophil counts. Further studies are needed to determine whether these genes affect immune cell function. Moreover, ssGSEA revealed differential enrichment of pathways associated with CD4^+^ T cells, myeloid cells, and dendritic cells. Because of the current lack of commercially available, validated canine-specific antibodies for FFPE samples, we were unable to perform IHC staining of these immune cell subsets. Further studies using appropriate antibodies will be required to validate these transcriptomic findings at the protein level.

This study had several limitations. First, it was a nonrandomized, open-label trial without a control arm. This may have limited the full assessment of treatment efficacy. Second, the cohort primarily consisted of dogs with advanced-stage disease, which may have resulted in the underestimation of the therapeutic effects of anti-PD-1 antibody during earlier stages. Third, although biomarker analyses were conducted, not all cases had sufficient samples for MSI testing, cytokine/chemokine profiling, IHC, or RNA-seq, which may have introduced selection bias and reduced statistical power. Fourth, the MSI analysis was conducted using a PCR-based method rather than whole-exome sequencing, and tumor TMB could not be assessed. Finally, because of the lack of validated antibodies for canine proteins, immunohistochemical validation of immune cell subsets identified by RNA-seq could not be performed. Similarly, PD-L1 expression in tumor cells could not be evaluated because a reliable anti-PD-L1 antibody for canine tissues was not available. These limitations highlight the need for follow-up studies with larger, stratified cohorts, standardized sample collection, and more advanced molecular profiling techniques.

In conclusion, the present study revealed that caninized anti-PD-1 antibody (ca-4F12-E6) is safe and shows clinical benefit in dogs with advanced OMM. Poor outcomes were associated with leukocytosis, neutrophilia, and elevated CRP, whereas MSI-high status predicted longer survival. These results support the use of canine OMM as a translational model for human studies.

## Declarations

### Ethics approval and consent to participate

#### Ethics approval

This study involved owned dogs and was approved by the Ethics Review Board of the Joint Faculty of Veterinary Medicine of Yamaguchi University and the Ethics Review Board of the Kyoto Animal Medical Center. Participants provided informed consent to the pet owners in the study before enrollment.

#### Patient consent for publication

Not applicable.

### Availability of data and material

Data are available upon reasonable request. Deidentified study data will be provided to other investigators for research purposes upon reasonable request.

### Authors’ contributions

Conceptualization, T.Mizuno. and M.I.; methodology, T.Mizuno. and M.I.; validation, T.K. and M.I.; formal analysis, T.Motegi.; investigation, M.I., K.H., S.I., H.M., K.I. and M.S.; resources, M.K. and T.T.; data curation, T.Mizuno. and T.K.; writing—original draft preparation, M.I. and T.Mizuno.; writing—review and editing, T.Motegi. and K.H.; visualization, M.I.; supervision, T.K. and T.Mizuno.; funding acquisition, T.Mizuno. All authors have read and agreed to the published version of the manuscript

## Acknowledgments

We thank Drs. Masaru Okuda, Satoshi Kambayashi, Kenji Baba, Munekazu Nakaichi, Hiroshi Sunahara, Harumichi Ito, Yuki Nemoto, and Kenji Tani for providing tumor samples. We are also grateful to the clinical staff at YUAMEC for their assistance in the clinics. Special thanks to Dr. Hidetoshi Uenaka (Real World Evidence Division, Pharmaceutical Business Unit, JMDC Inc.) for his support with statistical analysis.

---

**Figure S1.**
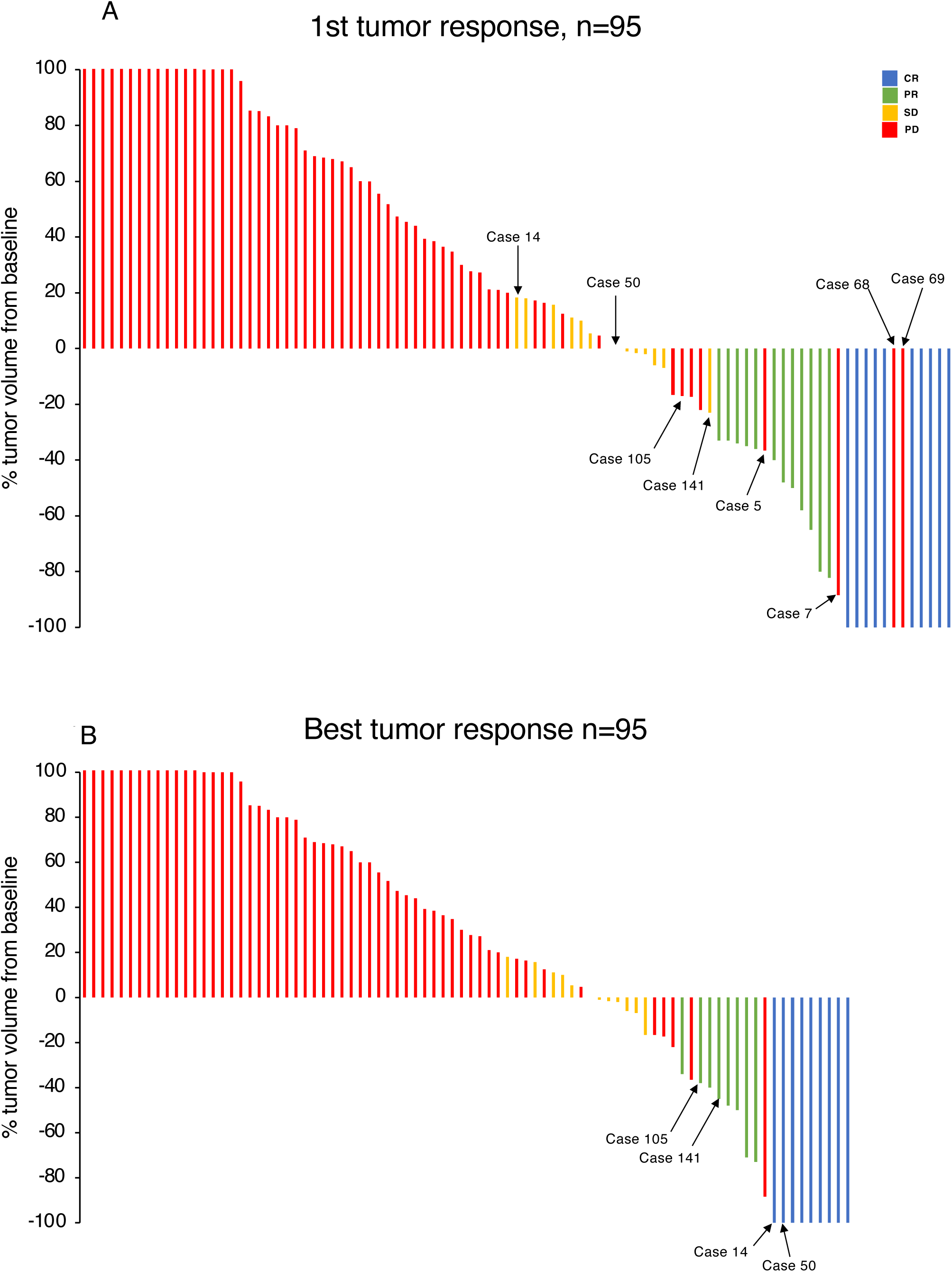
Changes in tumor volume following anti-PD-1 antibody treatment (A) Initial tumor response (n = 95) shown as the maximum percentage change in tumor diameter from baseline during the initial treatment period. (B) Best tumor response in each dog (n = 95) shown as the greatest percentage change in tumor diameter from the beginning of treatment to the end of the study. Tumor response categories included CR, PR, SD, and PD. The numbers indicated below each bar represent the case numbers referenced in the main text and correspond to the Case No. listed in Table S1.

**Figure S2.**
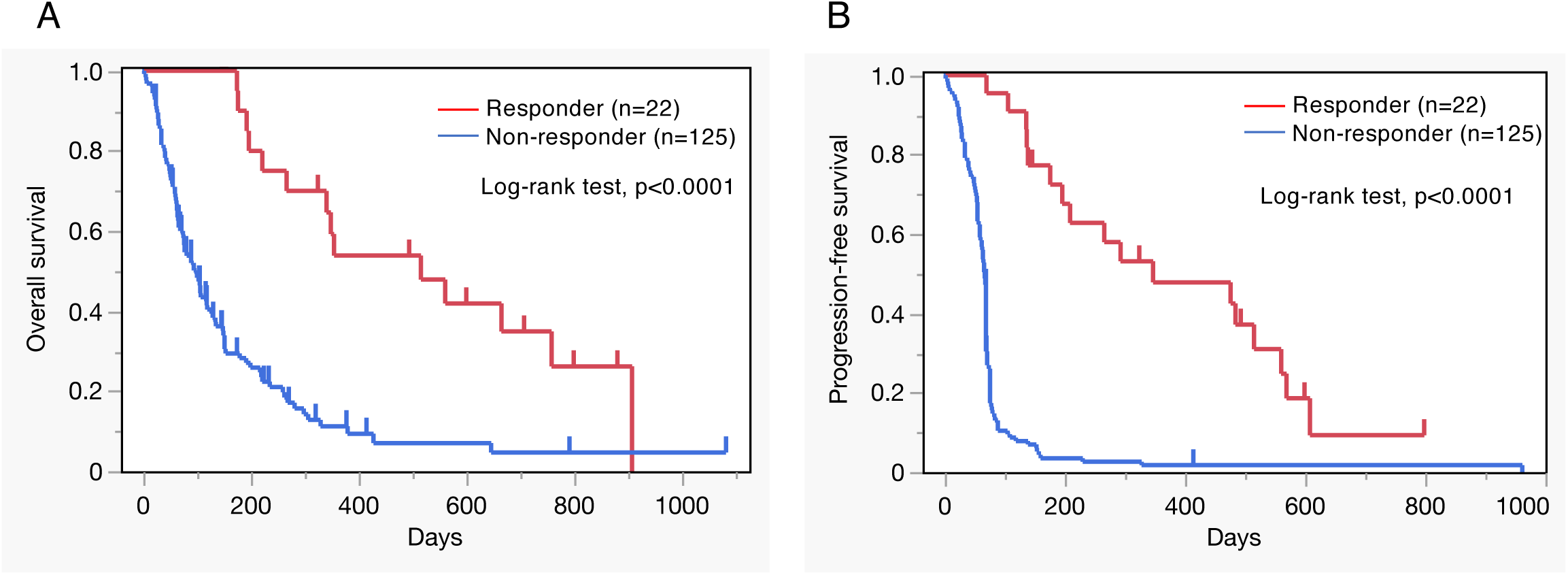
Survival analysis by treatment response (A) Kaplan–Meier curve for overall survival (OS) comparing responders (CR + PR, n = 22) and nonresponders (SD + PD, n = 125). Responders showed significantly longer OS (p < 0.0001). (B) Kaplan–Meier curve for progression-free survival (PFS) in the same groups. Responders also had significantly longer PFS (p < 0.0001).

